# Impact of inhibitors of histone post-translational modifications on lifespan, reproduction, and stress response in the rotifer *Brachionus manjavacas*

**DOI:** 10.1101/2025.05.03.652051

**Authors:** Nelia Luviano Aparicio, Meghan Dryburgh, Colleen M. McMaken, Alyssa Liguori, Kristin E. Gribble

## Abstract

Epigenetic modifications, including histone post-translational modifications, are central drivers of age-associated structural and functional changes in the genome, influencing gene expression and cellular resilience. Our objective was to determine the effects of inhibiting histone deacetylases (HDACs) and the histone methyltransferase SETDB1 on lifespan, reproduction, and stress response in the rotifer *Brachionus manjavacas*, a model organism for aging studies. We exposed rotifers to three pharmaceutical compounds, including the HDAC inhibitors β-hydroxybutyrate and sodium butyrate and the SETDB1 inhibitor mithramycin A. We quantified changes in the global histone modification levels by immunoblotting, and measured lifespan, reproduction, and heat stress resistance in the drug-treated rotifers relative to a control. Global histone acetylation levels increase with β-hydroxybutyrate and sodium butyrate treatments. Histone 3 K9 trimethylation (H3K9me3) levels were reduced by treatment with mithramycin A. β-hydroxybutyrate significantly extended lifespan without significantly modifying heat stress resistance. In contrast, mithramycin A increased lifespan and enhanced heat stress tolerance, demonstrating a dual protective effect. Sodium butyrate specifically improved heat stress resistance without affecting overall lifespan. Importantly, none of the three treatments had a significant impact on lifetime reproduction. These findings provide insights into the role of histone modifications in aging and suggest potential interventions targeting epigenetic marks to promote longevity and resilience.

## Introduction

Epigenetic mechanisms have emerged as key contributors to age-related changes in genome structure and function (Booth and Brunet 2016; López-Otín et al. 2023). Among the main types of epigenetic regulation, DNA methylation, histone modifications, and non-coding RNAs modify the architecture and accessibility of DNA and ultimately regulate gene expression (Alexander and Lomvardas 2014; Fraga and Esteller 2007; Gonzalo 2010; Oleksiak and Rajora 2020). In this study, we explore manipulation of epigenetic histone modifications as a means to increase lifespan and improve health.

Post-translational modifications (PTMs) of histone tails direct chromatin organization and gene expression through biochemical changes including acetylation, methylation, phosphorylation, ubiquitylation, and sumoylation. Specific enzymes catalyze the addition or removal of these chemical groups from lysine residues on histone tails, resulting in the formation of either condensed heterochromatin or accessible euchromatin. This dynamic restructuring of chromatin determines the accessibility of genomic regions to transcription factors, thereby precisely controlling gene expression patterns. These epigenetic modifications thus orchestrate essential physiological processes, from cell differentiation to organ development. The patterns of histone PTMs evolve throughout an organism’s lifespan, with acetylation and methylation emerging as the most extensively studied modifications in aging research (Y. Wang, Yuan, and Xie 2018).

Histone deacetylases (HDACs) remove acetyl groups from lysine residues on histones, leading to chromatin condensation and reduced transcriptional activity (Chang and Min 2002). Notably, different HDACs have distinct effects on lifespan. For example, deletion of the HDAC *SIR2* shortens the lifespan of *Saccharomyces cerevisiae*, while its overexpression extends lifespan (Kaeberlein, McVey, and Guarente 1999). In contrast, knocking out *RPD3*, another HDAC, significantly increases yeast lifespan (Kim, Villeponteau, and Jazwinski 1996). These contrasting effects illustrate the complexity of HDAC-mediated regulation of aging and emphasize the need to understand how specific HDACs influence longevity. Since aging is often associated with the downregulation or dysregulation of biosynthetic and metabolic pathways, treatment with HDAC inhibitors could potentially delay age-related decline by restoring expression of critical genes. Additionally, inhibiting HDACs may alter expression of genes involved in inflammation and stress responses, pathways closely linked to aging and longevity (Kourtis and Tavernarakis 2011; Pasyukova and Vaiserman 2017). These findings suggest that HDAC inhibitors hold promise as therapeutic tools for modulating aging processes.

The histone methyltransferase SETDB1 (SET domain bifurcated histone lysine methyltransferase 1, also known as ESET or KMT1E) catalyzes the deposition of di- and tri-methyl marks on H3K9 (H3K9me2 and H3K9me3), which are linked to transcriptional repression (Luo et al. 2023). Functional analysis and knockout studies have revealed that SETDB1 is necessary for establishing the methylation marks required for proper oocyte maturation and that *Setdb1* gene expression in mouse ovaries decreases with age (Bilmez, Talibova, and Ozturk 2022). This suggests that SETDB1 is involved in the alterations of H3K9 methylation patterns associated with aging, influencing gene expression and cellular function. Studies have reported a global change in H3K9me3 patterns with aging (Ocampo et al. 2024; Wood et al. 2010). In a mouse strain carrying a triple knockout of three methyltransferases responsible for H3K9me3 deposition, the loss of H3K9me3 in adulthood results in premature aging (Ocampo et al. 2024). A comparable phenomenon was observed in young *Drosophila melanogaster*, in which H3K9me3 is normally concentrated in specific chromosomal regions. As the flies age, this site-specific enrichment decreases, leading to similar levels of H3K9me3 across different chromosome regions (Wood et al. 2010).

Given that histone acetylation and methylation levels and distribution change with age and are associated with age-related changes in gene expression and functional decline, these epigenetic modifications are potential targets for manipulation to extend healthspan and lifespan. Pharmaceutical inhibitors of histone modifying enzymes, many of which have been used in treatment of cancer or other diseases, might be used as aging therapies, but relatively few have been tested in this capacity.

HDAC inhibitors prevent the removal of acetyl groups from histones, which can enhance gene expression and thereby possibly extend lifespan. In this work, we tested two HDAC inhibitors, β-hydroxybutyrate and sodium butyrate, that have been shown to increase lifespan in *Caenorhabditis elegans* (Edwards et al. 2014) and *Drosophila melanogaster* (Vaiserman et al. 2013), respectively. β-hydroxybutyrate is an endogenous ketone molecule that acts as a natural inhibitor of class I HDACs; it serves both as an energy source and as a signaling molecule that influences gene expression, inflammation, and stress resistance through epigenetic modifications (Shimazu et al. 2013). Sodium butyrate is a short-chain fatty acid that selectively inhibits class I and II HDACs, particularly targeting HDACs 1, 2, and 3. Its inhibitory effect is achieved by binding to the active site of HDAC enzymes, blocking their function and enhancing histone acetylation, which in turn promotes the transcription of genes involved in cell cycle regulation, apoptosis, and differentiation (Davie 2003).

Because of the association of high SETDB1 expression with reproductive aging in mice (Bilmez, Talibova, and Ozturk 2022), we additionally explored the lifespan and health effects of the SETDB1 inhibitor, mithramycin A. Mithramycin A is an antibiotic and anticancer compound known primarily for its ability to bind GC-rich DNA sequences, thereby inhibiting the transcription factor Sp1 and affecting gene expression (Miller et al. 1987). Recently, it has also been studied for its potential to inhibit histone methyltransferase SETDB1 (Federico et al. 2020). SETDB1 is crucial for gene silencing and maintaining heterochromatin structure, particularly in regions associated with oncogenesis and other disease states (Z. Zhao et al. 2022). By blocking SETDB1 activity, mithramycin A reduces H3K9me3 levels, leading to increased chromatin accessibility and the reactivation of silenced genes (Quarni et al. 2019).

To assess whether histone modification inhibitors could be used as therapies to increase longevity and improve health, we quantified age-specific gene expression of histone deacetylases and the histone methyltransferase SETDB1 and measured the effects of β-hydroxybutyrate, sodium butyrate, and mithramycin A on histone modification levels, lifespan, reproduction, and stress resistance in the rotifer *Brachionus manjavacas.* This microscopic, aquatic invertebrate is an effective experimental system in which to study aging due to its short life cycle, rapid generation time, ease of cultivation in the laboratory, and significantly higher genetic homology with humans than some other invertebrate laboratory model systems (Gribble & Mark Welch, 2017). Our findings will provide critical insights into how modulation of histone modifications can influence lifespan, reproduction, and stress resistance and potentially pave the way for novel therapeutic interventions to improve lifespan and late life health.

## Materials and methods

### Rotifer and algae culture

*Brachionus manjavacas* “L5 strain” was fed the chlorophyte *Tetraselmis suecica*, which was cultivated in 2 L flasks containing bubbled f/2 medium (Guillard 1975), excluding silica, prepared with 15 ppt Instant Ocean Sea Salt (Instant Ocean, Blacksburg, VA) made with filtered deionized water. Both the rotifer and algae cultures were maintained at 21°C under a 12:12 h light cycle. Cultures of *T. suecica* were kept in semi-continuous logarithmic growth phase by removing approximately 40-50% of the culture volume and replacing it with f/2 medium every two days throughout the experiments.

### Gene expression analysis

Age-specific gene expression of histone modifiers was collected as part of a larger RNA-seq study. Multiple generations of age synchronization were conducted before rotifers were collected. mRNA was extracted from five replicates of rotifer pellets from each of four different age cohorts: 1-, 3-, 6-, and 10-days old (∼400 rotifers per pellet for the 1-day timepoint, and ∼200 rotifers per pellet for later ages). RNA was extracted following the TRIzol™ reagent protocol (#15596026, Invitrogen), with residual DNA removed using Turbo DNA-free (#AM1907, Invitrogen), and RNA precipitated with isopropanol.

The quality, quantity, and purity of RNA were evaluated using an Agilent 2100 Bioanalyzer (Agilent Technologies), Qubit™ 2.0 Fluorometer (Invitrogen), and NanoDrop™ 2000 spectrophotometer (Thermo Fisher Scientific). cDNA libraries were constructed with poly-A enrichment from the RNA samples using the NEB Next Ultra II RNA kit for Illumina (#E7770, New England Biolabs). Sequencing was performed on an Illumina NovaSeq platform, generating 150 bp paired-end reads, through the eukaryotic mRNA sequencing service provided by Novogene Bioinformatics Technology.

The RNA-seq data were processed through a bioinformatics pipeline. Raw reads were trimmed using FASTX (Hannon 2010) and Cutadapt (Martin 2011) to remove adapter sequences and low-quality bases, ensuring that only high-quality reads were retained for downstream analysis. The cleaned reads were mapped to a reference genome using Bowtie2 (Langmead et al. 2019; Langmead and Salzberg 2012). The resulting FASTA files were subjected to blastn (Camacho et al. 2009) searches against the NCBI Nucleotide (nt) database. Sequences that yielded significant hits to non-metazoan organisms were excluded from the dataset. Transcript abundance was quantified using Salmon (Patro et al. 2017), which accounted for transcript length and sequencing depth to estimate the number of reads corresponding to each gene and gene isoform.

### Survival and reproduction assays

Lifespan and reproduction were measured as in previous studies (Bock et al. 2019; Liguori et al. 2024). To synchronize age and to control for maternal and grandmaternal ages of the experimental animals, two generations of maternal age synchronization were conducted prior to beginning life table experiments. Amictic eggs were removed from females by vortexing, isolated by micropipette, and allowed to hatch and mature for 5 days in *ad libitum* food conditions, at which time eggs were again collected from mature females. After repeating this for two generations, eggs were collected and hatched overnight to initiate the experimental cohort of same-aged individuals.

To initiate and track each cohort, individual neonates were deposited into 0.5 mL of *T. suecica* at a concentration of 6 × 10^5^ cells mL^−1^ in 15 ppt Instant Ocean and test drugs (as specified below) in 24-well plates (n = 72 per treatment). Individuals were observed every 24 h on a Zeiss Stemi 508 microscope, and survival, reproductive status (carrying or not carrying eggs), and the number of new offspring were quantified. The original female was then transferred to a well with new *T. suecica* and drug in 15 ppt Instant Ocean. Death was characterized by the cessation of swimming and of movement in the cilia, muscles, and mastax (jaw). Daily scoring and transfers were conducted until all individuals had died.

### Drug testing

β-hydroxybutyrate (#H6501, Sigma-Aldrich), sodium butyrate (#303410, Sigma-Aldrich) and mithramycin A (#11434, Cayman Chemical) were used to treat rotifers. Prior to conducting survivorship and reproduction assays, we tested 100 µM, 500 µM and 1 mM for β-hydroxybutyrate; 50 µM, 100 µM and 500 µM for sodium butyrate; and 100 nM, 500 nM and 1 µM for mithramycin A in a 3-day trial to determine optimal concentrations. Based on survivorship results, daily treatments for full experiments consisted of 1 mM β-hydroxybutyrate, 500 µM sodium butyrate, or 500 nM mithramycin A, which were added to experimental wells containing *T. suecica* in 15 ppt Instant Ocean. 0.01% DMSO was used as a control.

### Heat stress assay

To assess the impact of histone modification inhibitors on heat stress resistance, neonate rotifers were exposed to 1 mM β-hydroxybutyrate, 500 µM sodium butyrate, 500 nM mithramycin A, or 0.1% DMSO (control) for 2 days. Following treatment, 5 rotifers were placed in each of the 8 central wells of a 24-well plate, resulting in a total of 40 rotifers per treatment group. Each treatment was allocated to a separate plate. The plates were sealed with lab tape and floated in a 42 °C water bath for 1 hour to induce heat stress. After heat exposure, the plates were transferred to 21 °C. For each treatment, the number of live and dead rotifers was recorded at 24- and 48-hours post-heat exposure.

### Quantification of histone modifications

To assess whether histone modification inhibitors altered histone post-translational modifications (PTM) levels in *B. manjavacas*, three replicate 150 mL batch cultures were treated for three days with 500 µM β-hydroxybutyrate, 1 mM sodium butyrate, or 500 nM mithramycin A. After three days, rotifers were filtered onto a 100 µm sieve, transferred to 15 ppt Instant Ocean in a 6-well plate, and subjected to overnight starvation at 21°C to prepare the samples for histone extraction.

### Histone extractions

Histone extractions from *B. manjavacas* were performed using a hybrid protocol that combines elements of the acid extraction method (Shechter et al. 2007) with additional steps for subcellular fractionation, aimed at removing non-nuclear proteins more effectively. All steps were performed at 4 °C. Rotifers were pelleted via centrifugation at 10,000 rpm for 5 minutes. The rotifer pellet was resuspended in 500 µL of fresh, ice-cold hypotonic lysis buffer (HLB: 150 mM MgCl₂, 100 mM KCl, 1 M Tris-HCl [pH 8.0], 1M DTT, protease inhibitors, and nuclease-free water) and homogenized on ice using a Dounce homogenizer. The resulting homogenate was processed using a QIAshredder homogenizer (#79656, Qiagen) and centrifuged at 2,500 rpm for 10 minutes. The pellet was resuspended in the filtrate and incubated in 500 µL of fresh HLB for 30 minutes. Following centrifugation at 10,000 rpm for 10 minutes, the pellet was resuspended in 1 mL of fresh, ice-cold lysis buffer B (LBB: 750 mM NaCl, 1 M Tris-HCl [pH 8.0], 1% Ipegal, 1 M hexylene glycol, protease inhibitors, and nuclease-free water) and rotated at 4 °C for 30 minutes to further isolate nuclear components. After centrifugation at 8,500 rpm for 10 minutes, the pellet was resuspended in 400 µL of 0.4 N H₂SO₄, wrapped in parafilm, and rotated overnight to extract histones.

The samples were centrifuged at 13,000 rpm for 10 minutes to pellet debris, and the histone-containing supernatant was transferred to a new sterile tube. To precipitate the histones, 66 µL of trichloroacetic acid was added to the supernatant, followed by a 30-minute incubation on ice. The sample was centrifuged at 13,000 rpm for 10 minutes, and the resulting pellet was washed twice with 1 mL of ice-cold acetone. After the final wash, the acetone was removed, and the pellet was air-dried for 30 minutes. The dried histone pellet was resuspended in 50 µL of nuclease-free water and dissolved by heating at 95°C for 5 minutes. The histone concentration was determined using a Bradford protein assay (He 2011) to ensure adequate protein yield for downstream analyses.

### Western blot and dot blot analyses

After validation of the specificity and sensitivity of all antibodies targeting histone modifications and Histone 3 against rotifer histone protein extracts by Western blot (Supplementary file 1, Fig S1), we proceeded with the dot blot analyses (Supplementary file 1, Fig S2). For both assays, polyvinylidene fluoride (PVDF) membranes were used for protein transfer and binding and antibody incubations.

For dot blots, histone extracts from controls, and from β-hydroxybutyrate, sodium butyrate, and mithramycin A-treated samples were analyzed. Three replicates of each sample were normalized to 700 ng of total protein. As a positive control, histones from sodium butyrate-treated HeLa cells (#3620, Active Motif) were included to validate the assay’s sensitivity to histone acetylation. An unmodified histone H3 peptide (#ab7228, Abcam) served as a negative control, ensuring specificity for modified histones. Denatured samples (2 µL each) were spotted in triplicate onto a methanol-activated PVDF membrane, followed by UV crosslinking using a Stratalinker® UV Crosslinker (Autocrosslink setting).

For Western blots, two replicates for each of controls, β-hydroxybutyrate-treated, and mithramycin A-treated samples were separated by SDS-PAGE on replicate gels loaded with the same samples and concentrations. Histone extracts were 2µg/µL for all treatments. Gels were transferred overnight onto PVDF membranes using a wet-transfer apparatus at 40 V.

For Western and dot blots, membranes were blocked with TBST (1× TBS 0.05% Tween 20) and 5% powdered milk for 1 hour at 37°C under elliptical agitation. Each membrane was treated separately with a different primary antibody (1:1000 dilution in blocking solution) and incubated overnight. The primary antibodies used include anti-histone H3 (#24834, Abcam), anti-histone H3 pan-acetyl (#ab300641, Abcam), or anti-histone H3K9me3 (#A-4036, Epigentek).

After primary antibody incubation, the membranes were washed for 10 minutes three times with TBST under elliptical agitation and subsequently incubated for 1 hour at room temperature in a 1:10000 dilution of secondary antibody. HRP-conjugated Goat anti-Rabbit IgG (H+L) secondary antibody (#AS014, ABClonal) was used for the membranes previously incubated with anti-Histone 3 pan-acetyl or anti-H3K9me3. HRP-conjugated Goat anti-Mouse IgG (H+L) (#AS003, ABClonal) was used for the membrane previously incubated with anti-Histone 3. The membranes were washed again three times with TBST and developed using SuperSignal™ West Pico chemiluminescent substrate (#34096, Thermo Fisher Scientific). Luminescence was imaged using an Amersham^TM^ Imager 600 with a 60-second exposure.

### Statistical Analyses

Statistical analyses were performed in RStudio using several specialized libraries to ensure robust data handling, visualization, and hypothesis testing. The “tidyverse” (Hadley Wickham et al. 2019) package was used for data manipulation and visualization, incorporating functions from “ggplot2” (H Wickham et al. 2016) for creating graphics and “dplyr” (Hadley Wickham et al. 2014) for efficient data wrangling. After verifying the normality of the data distribution with the Shapiro-Wilk test and testing for homogeneity of variances with the Levene’s test for the histone modification levels, Welch’s t-tests were used to compare histone modification levels between the treatments and controls.

Kaplan-Meier survival curves were generated to estimate the probability of survival over time for each treatment group (n = 72 per group) with the “survival” package (Therneau, 2015). Median survival times with 95% confidence intervals were calculated. The log-rank (Mantel-Cox) test was used to assess differences in survival distributions between treatment groups. Pairwise comparisons between treatment groups were conducted using the log-rank test with Benjamini-Hochberg correction for multiple comparisons. Survivorship curves were visualized using the “survminer” package (Pawar, Chowdhury, and Salvi 2022). Statistical significance was set at *p* < 0.05.

To assess differences in reproductive output across treatments, initial normality and homogeneity of variance were tested using Shapiro-Wilk and Bartlett’s tests. Given that the data violated parametric assumptions (*p* < 0.05 for both tests), we conducted Kruskal-Wallis tests to evaluate differences in reproductive metrics (lifetime reproductive output, total non-viable eggs per individual and reproductive period) among treatments. When the Kruskal-Wallis test revealed significant differences (*p* < 0.05), we performed post-hoc pairwise comparisons using Wilcoxon rank-sum tests with Bonferroni correction to control for multiple comparisons.

Non-parametric comparisons between groups exposed to heat-stress were performed using the Friedman test, followed by Wilcoxon signed-rank post-hoc tests to compare each treatment against the controls with the “coin” package (Hothorn et al. 2008).

## Results

### Gene expression and histone modifications

We found that HDACs and SETDB1 enzymes are differentially expressed across age in *B. manjavacas*. Expression of HDACs and SETDB1 was lowest in early life (1 day old) and increased through mid-life reaching peak expression at 10 days old (Fig 1, Supplementary file 2, Table S1).

**Fig 1.**
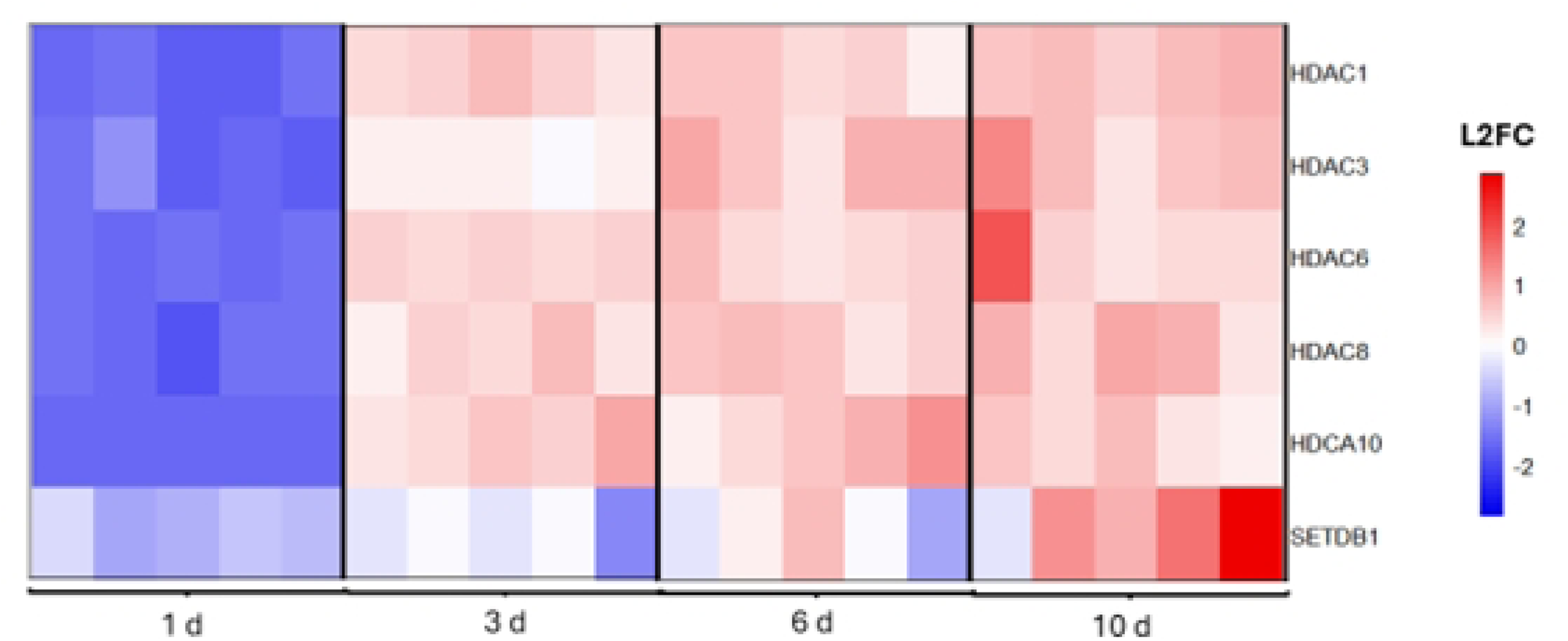
Relative gene expression levels of histone deacetylase enzymes (HDACs) and the SETDB1 enzyme in the rotifer *Brachionus manjavacas* across lifespan (1-, 3-, 6-, and 10-days old). Each age group includes five replicates. L2FC stands for Log2 Fold Change.

Dot blot analyses were conducted to quantify changes in histone modification levels in rotifers following treatment with β-hydroxybutyrate, sodium butyrate, or mithramycin A (Fig 2). Histone acetylation levels increased significantly in response to treatment with the deacetylation inhibitors β-hydroxybutyrate (*t* = 3.38, *df* = 8, *p* = 0.009) and sodium butyrate (*t* = 4.65, *df* = 8, *p* < 0.001), indicating enhanced acetylation activity compared to controls (Fig 2A and 2C). Mithramycin A treatment led to a significant reduction in H3K9me3 levels (*t* = 2.38, *df* = 16, *p* = 0.02) (Fig 2E).

**Fig 2.**
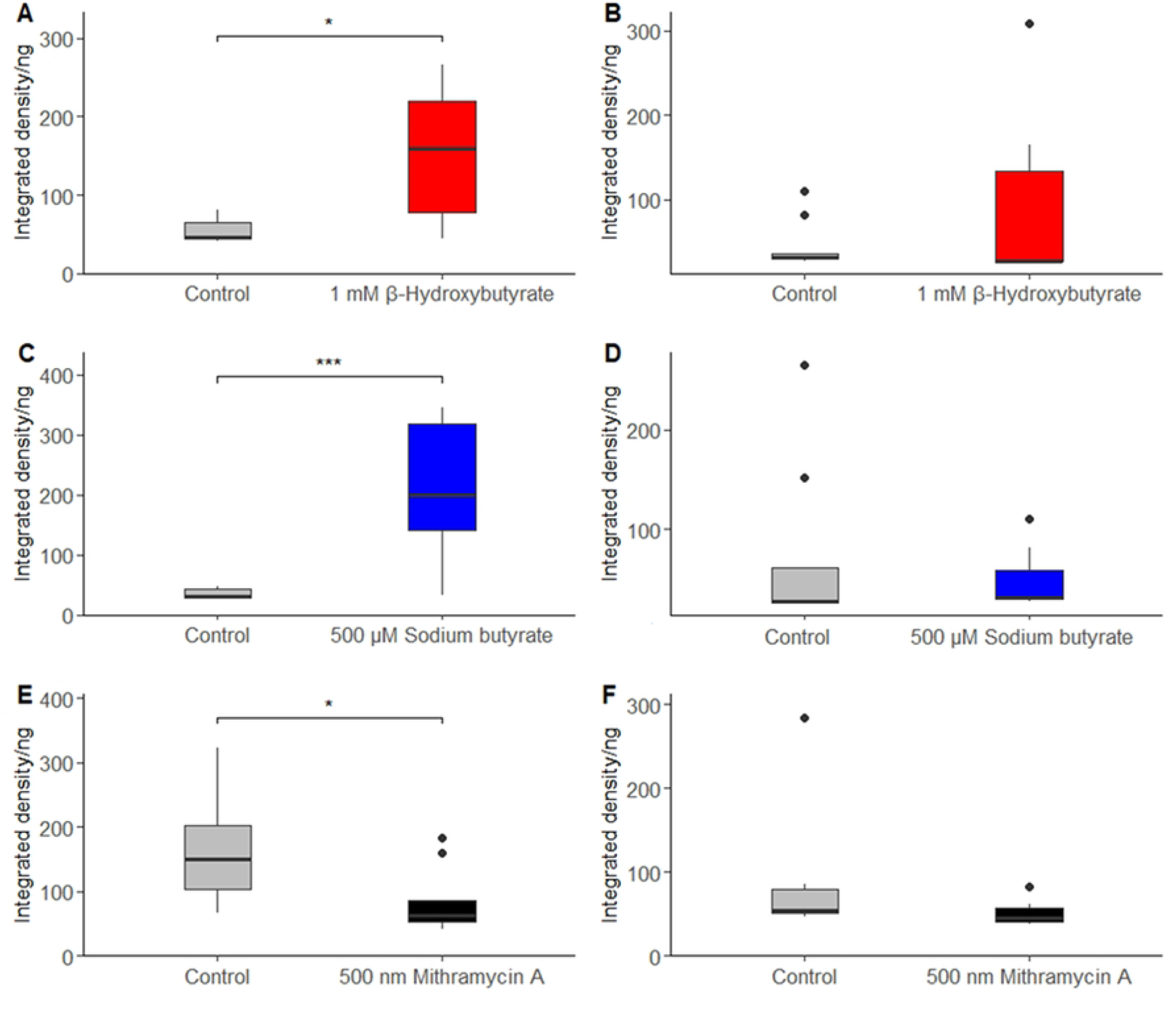
Quantification of histone modifications and histone H3 levels in *B. manjavacas* following drug treatment, measured by dot blot analysis. The horizontal bars show the median values, with the boxes representing the interquartile range (IQR) from the 25th to 75th percentile, and error bars denoting range. Histone modification levels: histone 3 acetylation in response to β-hydroxybutyrate (A) and sodium butyrate (C), as well as H3K9me3 following treatment with mithramycin A (E). Corresponding histone H3 levels after treatment with β-hydroxybutyrate (B), sodium butyrate (D), and mithramycin A (F). Statistical significance between control and treated groups is denoted by asterisks, where * indicates p < 0.05 and *** indicates p < 0.001.

We also examined histone H3 levels across all treatments. While β-hydroxybutyrate and sodium butyrate specifically impacted histone acetylation and mithramycin A affects H3K9 methylation, none of these treatments significantly alter the total histone H3 content in *B. manjavacas* (Fig 2B, 2D, and 2F). These data collectively suggest that the observed changes in epigenetic modifications levels are due specifically to alterations in post-translational modifications and not to changes in histone H3 abundance.

### Lifespan and reproduction effects of histone modification inhibitors

β-hydroxybutyrate and mithramycin A both significantly extend median and maximum lifespan relative to the control. Sodium butyrate exhibits a survivorship trend similar to that of the control group (Fig 3). Median lifespan was 11 days for the control and sodium butyrate-treated groups. β-hydroxybutyrate and mithramycin A treatments significantly increased median lifespan to 14 days (pairwise log-rank tests, p=0.030 and p=0.006, respectively). Similar trends were seen for maximum lifespan (5% survivorship) which was 19 d for the control and sodium butyrate-treated groups and increased to 21 d for β-hydroxybutyrate and mithramycin A treated groups (Fig 3, Supplementary file 2, Table S2). Conversely, reproductive measures were not significantly affected by any of the drug treatments (Fig 4, Supplementary file 2, Table S3).

**Fig 3.**
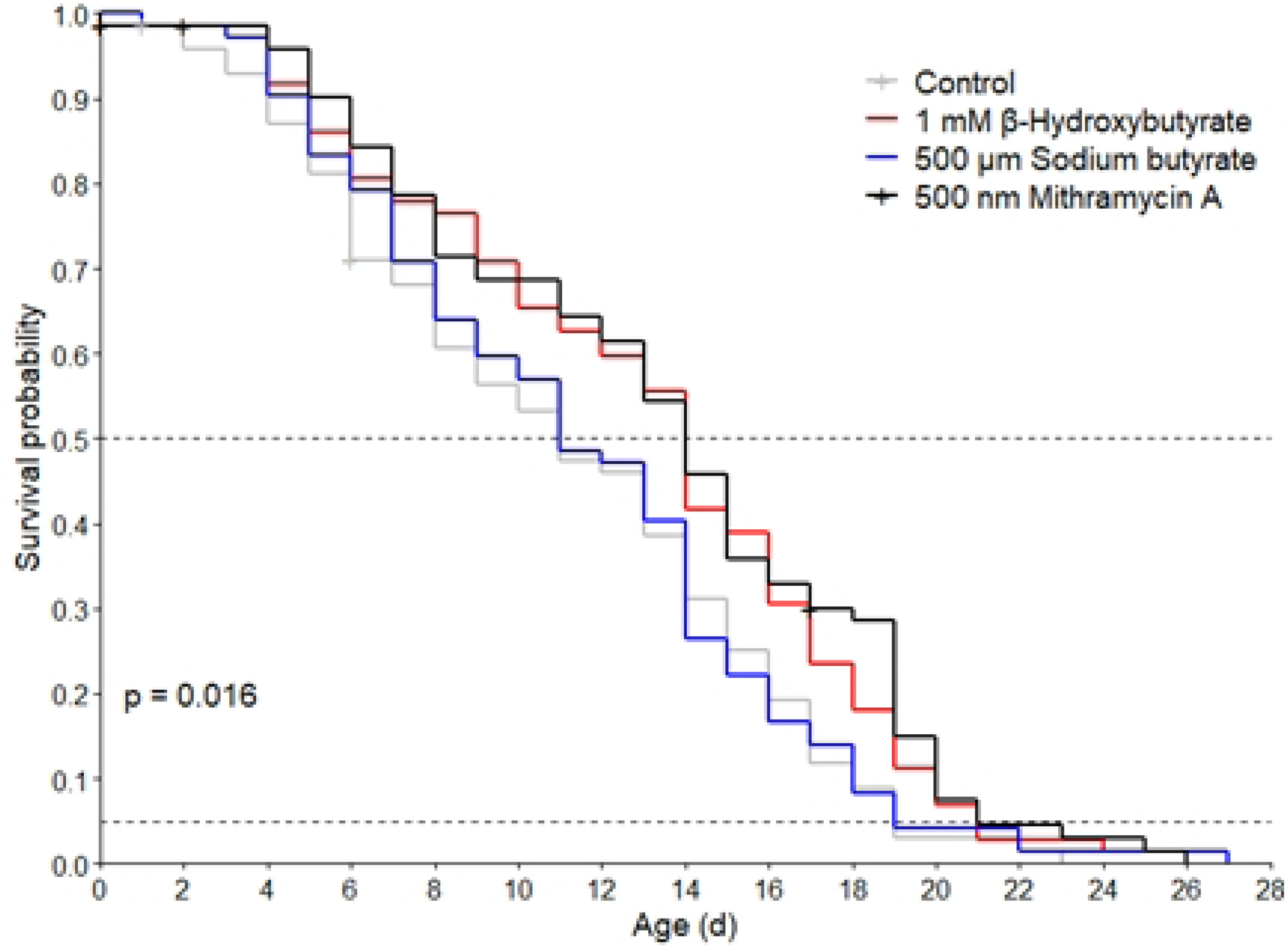
Survivorship of *B. manjavacas* in control group (grey) and under three drug treatments: β-hydroxybutyrate (red), sodium butyrate (blue) and mithramycin A (black). The *p*-value represents the Mantel Cox test across all 4 survivorship curves. Dashed lines indicate and median (50% survivorship) and maximum (5% survivorship) lifespan.

**Fig 4.**
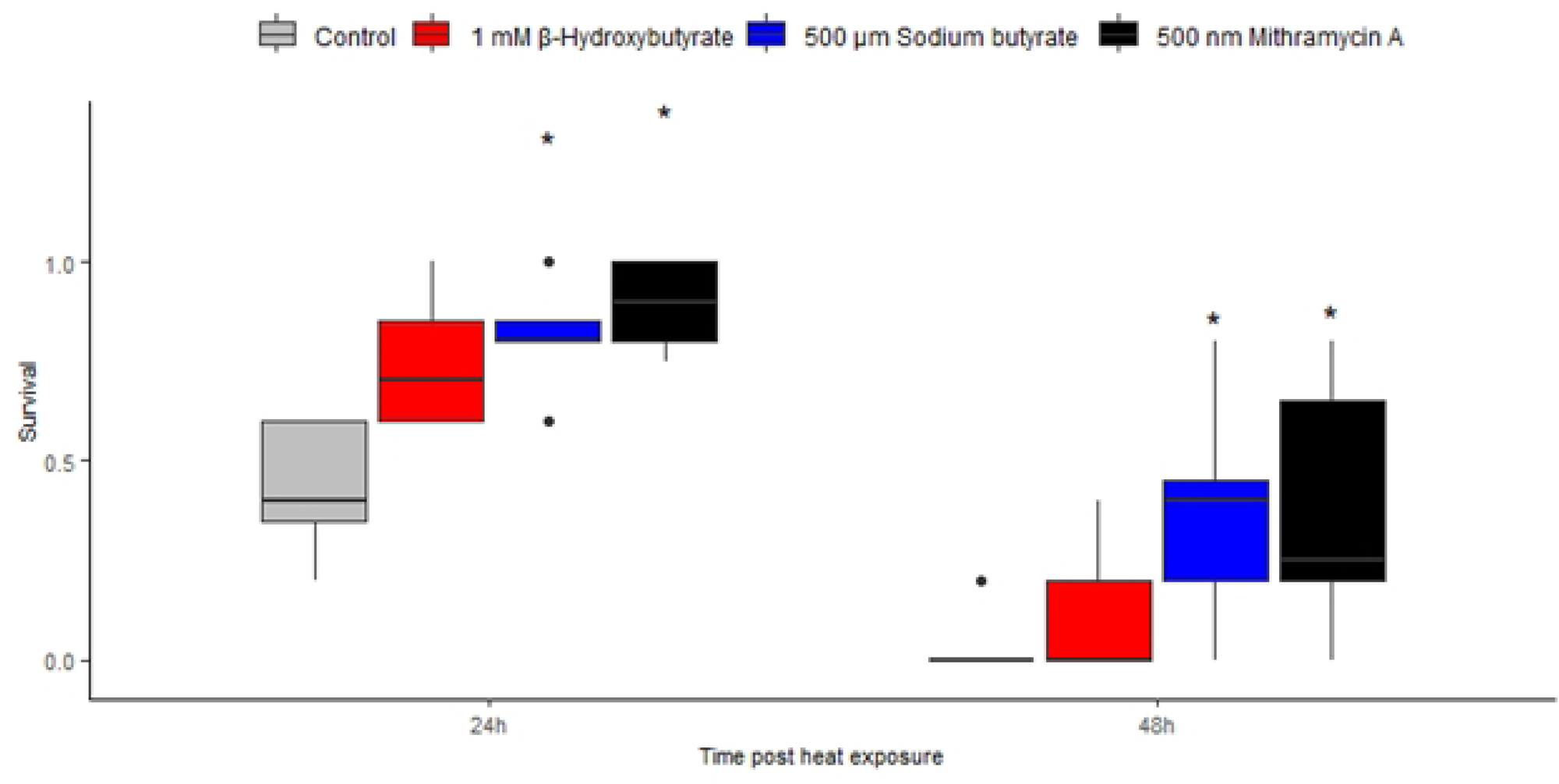
Measures of reproduction in *B. manjavacas* across control and treatment groups: β-hydroxybutyrate (BHB), sodium butyrate (SB), and mithramycin (MTA): (A) Lifetime reproductive output (offspring per individual), (B) daily reproduction per individual, (C) total non-viable eggs per individual, and (D) reproductive period (in days). The bars show the median values, with the error bars representing the interquartile range from the 25th to 75th percentile.

### Effect of histone modification inhibitors on the heat stress resistance

Histone modification inhibitor treatments significantly enhanced rotifer survival relative to that of the control at both 24- and 48-hours following heat exposure (Friedman test, maxT = 4.64, p < 0.0001). Specifically, sodium butyrate and mithramycin A markedly increased the rotifers’ ability to withstand elevated temperatures at both time points, with survival rates significantly higher than the control group (p < 0.005 for each). In contrast, β-hydroxybutyrate exhibited a more modest protective effect, resulting in survival improvements that approached but did not achieve statistical significance (p = 0.08) (Fig 5, Supplementary file 2, Table S4).

**Fig 5.**
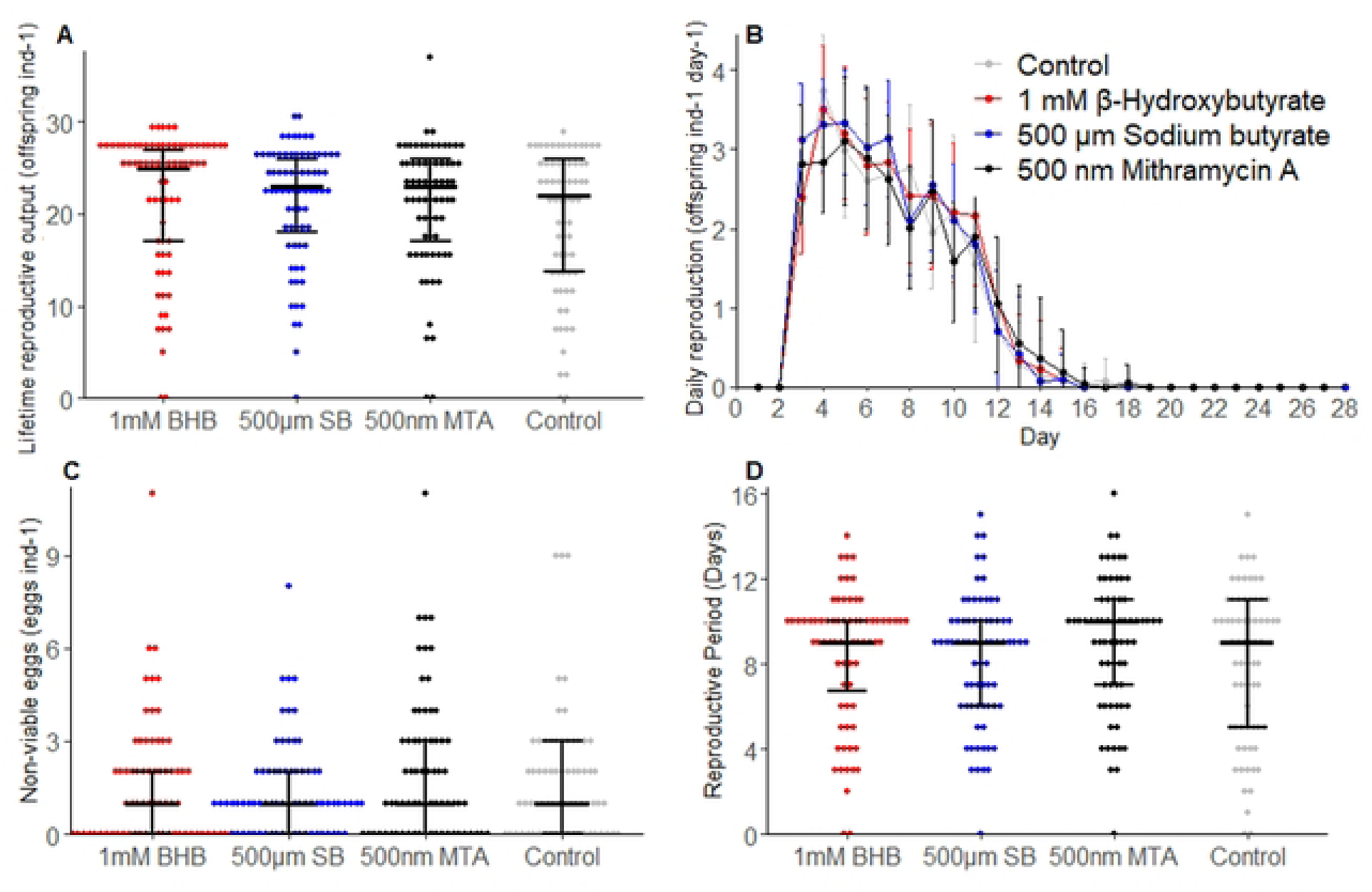
Heat stress resistance of *B. manjavacas* at 24- and 48-hours after heat exposure. The horizontal bars in the plots show the median values, with the boxes representing the interquartile range (IQR) from the 25th to the 75th percentile, and error bars denoting range. Statistical significance between control and treated groups is denoted by asterisks, where * indicates p < 0.05.

## Discussion

Epigenetic modifications are increasingly being shown to be important as controls on health and lifespan rather than simply marks correlated with biological age (Izadi et al. 2024; K. Wang et al. 2022). The reversible nature of epigenetic marks makes them promising targets to modulate aging rate and phenotype. In this study, we investigated the use of histone modification inhibitors as a means to extend lifespan and increase stress resistance. We found that gene expression of HDACs 3, 5, 6, and 8, and of the histone methyltransferase SETDB1, increases with increasing age in *B. manjavacas*. A prior transcriptomic analysis of aging in *B. manjavacas* showed a similar age-related increase in expression of HDACs 1, 3, 4, 6, and 8, highlighting the connection of HDACs to aging in rotifers (Gribble and Mark Welch 2017). The previous study sampled at different, and less specific ages: eggs, neonates (1–3 hours old), early reproductive females (36 hours old), mixed-age reproductive females (3–6 days old), and post-reproductive females (6–9 days old). These age-related increases in HDAC and SETDB1 expression would be expected to result in decreased levels of histone acetylation and increased histone methylation, leading to a more condensed chromatin structure and reduced gene transcription (Bannister and Kouzarides 2011).

Age-related changes in the expression of histone-modifying enzymes, such as histone deacetylases (HDACs) and histone methyltransferases, have been observed in some vertebrate species and have been shown to impact chromatin structure and gene regulation. Relative expression of HDACs increases with age in human cerebral white matter and is associated with age-related changes in its structure. A post-mortem study further confirmed that HDAC 1 and HDAC 2 paralogs are significantly elevated in white matter tissue from elderly individuals (Gilbert et al. 2019). High levels of HDACs are also detected in aged mice and human brain samples. HDAC 1, 2, 3, 6 and 7 were found to be specifically intense in the hippocampal formation from adult mice, while a significant elevation of HDAC 1, 3 and 7 were found in the hippocampus in aged human brain samples (Auzmendi-Iriarte et al. 2022). In the short-lived fish *Nothobranchius furzeri*, HDAC 1 expression decreases with age in muscle, liver, and brain tissues (Zupkovitz et al. 2018).

Less information about the age-related expression of HDACs 1-8 is available for invertebrate models. Knockdown experiments targeting histone deacetylases (HDACs) have been conducted in *Drosophila melanogaster* to explore their roles in aging. Specifically, the knockdown of HDAC 6 in flies led to significant extensions in both lifespan and healthspan, with the most pronounced effects observed when HDAC 6 was silenced specifically in neuronal tissues (Y. Zhao et al. 2022). These findings suggest that age-associated alterations in histone-modifying enzymes like HDAC 6 are conserved across species and contribute to aging through epigenetic regulation.

The age-associated increases in histone deacetylation and H3K9me3 that accompany changes in expression of HDACs and SETDB1 led us to hypothesize that decreasing the activity of these enzymes could change lifespan and healthspan outcomes. Thus, we pharmacologically altered histone acetylation and methylation using drugs that inhibit histone modifiers. We found a significant increase in histone acetylation levels in response to the deacetylation inhibitors β-hydroxybutyrate and sodium butyrate, indicating reduced deacetylation activity compared to controls. This increase suggests that both β-hydroxybutyrate and sodium butyrate treatments promote a more open chromatin state conducive to increased gene expression.

Treatment with β-hydroxybutyrate extends the lifespan of *B. manjavacas* compared to untreated controls. These findings in rotifers are consistent with studies in *Caenorhabditis elegans*, where β-hydroxybutyrate extends lifespan in a concentration-dependent manner (Edwards et al. 2014). β-hydroxybutyrate also increased the mean survival of worms by 22% at elevated temperatures, suggesting enhanced thermotolerance. However, no induction of heat shock protein expression was observed in four heat shock reporter strains (*hsp-6::gfp*, *hsp-60::gfp*, *hsp-4::gfp*, and *hsp-16.2::gfp*), suggesting that while β-hydroxybutyrate improves thermotolerance, it does not activate a generalized heat shock response. In our study, β-hydroxybutyrate slightly improved heat stress resistance in *B. manjavacas.* This effect may be attributed to increased histone acetylation that promotes the expression of stress response genes that help mitigate heat-induced cellular damage (Horowitz 2016).

Sodium butyrate did not significantly extend the median lifespan of *B. manjavacas*. Our results differ from those in *D. melanogaster*, where the impact of sodium butyrate on lifespan varies depending on the timing of administration. Specifically, when administered during senescent stages, sodium butyrate reduces mortality rates and increases longevity, whereas administration exclusively during early life or throughout the entire adult lifespan decreases longevity (McDonald, Maizi, and Arking 2013). These complex, stage-specific effects align with the nuanced roles that histone acetylation plays in aging and stress resistance across species. It is possible that administering sodium butyrate only in late life in rotifers could yield different effects on lifespan. In our study, sodium butyrate had no significant impact on reproductive output in *B. manjavacas*, but it did enhance heat stress resistance. This suggests that its primary benefits may relate more to stress resilience rather than to reproduction or lifespan, likely due to its role in maintaining histone acetylation levels that modulate stress-responsive gene expression (Xu et al. 2022). Histone 3 acetylation may thus be a key regulatory mechanism in the adaptive response to thermal stress.

Mithramycin A has been widely studied for its ability to bind DNA and modulate gene expression, particularly in the context of cancer and neurological research (Z. Zhao et al. 2022; Voisine et al. 2007). Most studies have focused on its effects in mice and in cell culture systems (Geng et al. 2021). Given its role as a transcriptional inhibitor specifically targeting Sp1 transcription factors (Miller et al. 1987), mithramycin A has the potential to be used to investigate epigenetic modifications and gene regulation in aging. We tested mithramycin A for new uses in aging using rotifers, given that research on mithramycin A in common invertebrate models (e.g., *C. elegans*) is limited (Voisine et al. 2007). Mithramycin A treatment led to a significant reduction in H3K9me3 levels in *B. manjavacas,* likely causing a decrease in repressive methylation marks, which may allow reactivation of previously silenced genes. Mithramycin A significantly extended median lifespan of *B. manjavacas,* showing an effect on longevity comparable to that of β-hydroxybutyrate. This effect may be linked to its inhibition of the transcription factor Sp1, which is known to interact with SETDB1, affecting gene expression patterns relevant to aging and cellular stress responses (Miller et al. 1987). Unlike the other treatments, mithramycin A increased reproductive period slightly. This improvement could be related to its inhibition of SETDB1, which alters H3K9me3 patterns, potentially influencing gene expression linked to reproduction (Bilmez, Talibova, and Ozturk 2022). While the enhancement was not statistically significant, the observed trend warrants further investigation to understand the nuanced role of SETDB1 inhibition on reproduction. Furthermore, mithramycin A was shown to enhance the heat stress resistance of rotifers, suggesting that the observed reduction in H3K9me3 regulates key genes involved in heat stress adaptation. This observation is consistent with findings in plant species, which indicate that histone methylation, specifically histone 3 K4 dimethylation, is essential for regulating thermotolerance by influencing the transcription of heat stress-related genes (S. He et al. 2022). Collectively, these results highlight that alterations in histone modifications are fundamental mechanisms through which various organisms can adapt to thermal stress.

None of the drugs tested decreased lifetime reproductive output or increased the number of non-viable embryos significantly, suggesting that the observed improvements in lifespan and stress resistance were not due to energy or resource trade-offs with fecundity. The results further suggest that the global changes in histone deacetylation and H3K27me3 levels caused by the tested compounds do not negatively affect development in *B. manjavacas*.

## Conclusions

This study demonstrates the important role that histone modifications play in regulating lifespan and stress response and emphasizes the potential of HDAC and SETDB1 inhibitors as therapeutic agents to promote longevity and improve health. How the global changes in histone acetylation and H3K27me3 levels induced by the drugs tested here lead to beneficial outcomes remains to be determined. We hypothesize that β-hydroxybutyrate, sodium butyrate, and mithramycin A act by upregulating the expression of genes that are normally down-regulated in late age. Those genes that typically maintain high late-life expression presumably are not further upregulated by opening of chromatin at additional sites, and thus would likely not be impacted by drug treatment. Future research should explore the intergenerational and transgenerational consequences of these compounds, as well as the effects of additional histone modification inhibitors, to elucidate the molecular mechanisms underpinning their impact on aging.

## CRediT authorship contribution statement

**Kristin E. Gribble:** Writing – Review & Editing, Validation, Supervision, Funding Acquisition, Resources, Project administration, Methodology, Investigation, Conceptualization. **Nelia Luviano Aparicio:** Writing – Original Draft, Formal analysis, Data curation, Conceptualization, Investigation. **Colleen M. McMaken:** Writing – Review & Editing, Methodology, Formal analysis, Data curation. **Meghan Dryburgh:** Methodology, Formal analysis, Data curation. **Alyssa Liguori:** Writing – Review & Editing, Investigation, Methodology, Validation, Visualization.

## Financial Disclosure Statement

This project was funded by NSF CAREER Award IOS-1942606 to KEG. MD was funded by the Washington and Jefferson University Internship to the MBL. The funders had no role in study design, data collection and analysis, decision to publish, or preparation of the manuscript

## Competing interests

The authors declare that they have no conflict of interest.

## Acknowledgements

The authors thank Lucas Pender for laboratory assistance, Sovannarith Korm for guidance in the Western blotting protocol, and Emily Stone for assistance in developing the histone extraction protocol for *B. manjavacas*.

## Data availability

Western Blot and dot blot images, and data for survivorship, reproduction, heat stress resistance, and gene expression are provided as supplementary files for this manuscript.

## Notes

### Competing Interest Statement

The authors have declared no competing interest.

## References

1. Alexander JM, Lomvardas S. Nuclear architecture as an epigenetic regulator of neural development and function. Neuroscience. 2014;264:39–50.

2. Auzmendi-Iriarte J, Moreno-Cugnon L, Saenz-Antoñanzas A, Grassi D, de Pancorbo MM, Arevalo M-A, et al. High levels of HDAC expression correlate with microglial aging. Expert Opinion on Therapeutic Targets. 2022;26(10):911–22.

3. Bannister AJ, Kouzarides T. Regulation of chromatin by histone modifications. Cell research. 2011;21(3):381–95.

4. Bilmez Y, Talibova G, Ozturk S. Expression of the histone lysine methyltransferases SETD1B, SETDB1, SETD2, and CFP1 exhibits significant changes in the oocytes and granulosa cells of aged mouse ovaries. Histochemistry and Cell Biology. 2022;158(1):79–95.

5. Bock MJ, Jarvis GC, Corey EL, Stone EE, Gribble KE. Maternal age alters offspring lifespan, fitness, and lifespan extension under caloric restriction. Scientific Reports. 2019;9(1):3138.

6. Booth LN, Brunet A. The aging epigenome. Molecular Cell. 2016;62(5):728–44.

7. Camacho C, Coulouris G, Avagyan V, Ma N, Papadopoulos J, Bealer K, et al. BLAST+: architecture and applications. BMC Bioinformatics. 2009;10:421.

8. Chang KT, Min K-T. Regulation of lifespan by histone deacetylase. Ageing Research Reviews. 2002;1(3):313–26.

9. Davie JR. Inhibition of histone deacetylase activity by butyrate. The Journal of Nutrition. 2003;133(7):2485S–93S.

10. Edwards C, Canfield J, Copes N, Rehan M, Lipps D, Bradshaw PC. D-beta-hydroxybutyrate extends lifespan in C. elegans. Aging (Albany NY). 2014;6(8):621.

11. Federico A, Steinfass T, Larribère L, Novak D, Morís F, Núñez L-E, et al. Mithramycin A and mithralog EC-8042 inhibit SETDB1 expression and its oncogenic activity in malignant melanoma. Molecular Therapy-Oncolytics. 2020;18:83–99.

12. Fraga MF, Esteller M. Epigenetics and aging: the targets and the marks. Trends in Genetics. 2007;23(8):413–8.

13. Geng H, Su Y, Huang R, Fan M, Li X, Lu X, Sheng H. Specific protein 1 inhibitor mithramycin A protects cardiomyocytes from myocardial infarction via interacting with PARP. In Vitro Cellular & Developmental Biology-Animal. 2021 Mar;57:315–23.

14. Gilbert TM, Zürcher NR, Catanese MC, Tseng C-EJ, Di Biase MA, Lyall AE, et al. Neuroepigenetic signatures of age and sex in the living human brain. Nature Communications. 2019;10(1):2945.

15. Gonzalo S. Epigenetic alterations in aging. Journal of Applied Physiology. 2010;109(2):586–97.

16. Gribble KE, Mark Welch DB. Genome-wide transcriptomics of aging in the rotifer *Brachionus manjavacas*, an emerging model system. BMC Genomics. 2017;18:1–14.

17. Guillard RR. Culture of phytoplankton for feeding marine invertebrates. In: Culture of marine invertebrate animals: Proceedings—1st conference on culture of marine invertebrate animals Greenport 1975 Oct (pp. 29-60). Boston, MA: Springer US.

18. Hannon GJ. FASTX-Toolkit: FASTQ/A short-reads pre-processing tools. Repository http://hannonlabcshledu/fastx_toolkit. 2010.

19. He F. Bradford protein assay. Bio-Protocol. 2011:e45-e.

20. He S, Zhang Y, Wang J, Wang Y, Ji F, Sun L, et al. H3K4me2, H4K5ac and DNA methylation function in short-and long-term heat stress responses through affecting the expression of the stress-related genes in *G. hirsutum*. Environmental and Experimental Botany. 2022;194:104699.

21. Horowitz M. Epigenetics and cytoprotection with heat acclimation. Journal of Applied Physiology. 2016;120(6):702–10.

22. Hothorn T, Hornik K, Van De Wiel MA, Zeileis A. Implementing a class of permutation tests: the coin package. Journal of Statistical Software. 2008;28(8):1–23.

23. Izadi M, Sadri N, Abdi A, Serajian S, Jalalei D, Tahmasebi S. Epigenetic biomarkers in aging and longevity: Current and future application. Life Sciences. 2024:122842.

24. Kaeberlein M, McVey M, Guarente L. The SIR2/3/4 complex and SIR2 alone promote longevity in *Saccharomyces cerevisiae* by two different mechanisms. Genes & Development. 1999;13(19):2570–80.

25. Kim S, Villeponteau B, Jazwinski SM. Effect of replicative age on transcriptional silencing near telomeres inSaccharomyces cerevisiae. Biochemical and Biophysical Research Communications. 1996;219(2):370–6.

26. Kourtis N, Tavernarakis N. Cellular stress response pathways and ageing: intricate molecular relationships. The EMBO Journal. 2011;30(13):2520–31.

27. Langmead B, Salzberg SL. Fast gapped-read alignment with Bowtie 2. Nature Methods. 2012;9(4):357–9.

28. Langmead B, Wilks C, Antonescu V, Charles R. Scaling read aligners to hundreds of threads on general-purpose processors. Bioinformatics. 2019;35(3):421–32.

29. Liguori A, Korm S, Profetto A, Richters E, Gribble KE. Maternal age effects on offspring lifespan and reproduction vary within a species. Ecology and Evolution. 2024;14(5):e11287.

30. López-Otín C, Blasco MA, Partridge L, Serrano M, Kroemer G. Hallmarks of aging: An expanding universe. Cell. 2023;186(2):243–78.

31. Luo H, Wu X, Zhu XH, Yi X, Du D, Jiang DS. The functions of SET domain bifurcated histone lysine methyltransferase 1 (SETDB1) in biological process and disease. Epigenetics & Chromatin. 2023;16(1):47.

32. Martin M. Cutadapt removes adapter sequences from high-throughput sequencing reads. EMBnet Journal. 2011;17(1):10–2.

33. McDonald P, Maizi BM, Arking R. Chemical regulation of mid-and late-life longevities in *Drosophila*. Experimental Gerontology. 2013;48(2):240–9.

34. Miller DM, Thomas SD, Ray R, Sanchez J, Polansky DA, Campbell VW, et al. Rapid communication: Mithramycin selectively inhibits transcription of GC containing DNA. The American Journal of the Medical Sciences. 1987;294(5):388–94.

35. Ocampo A, Mrabti C, Yang N, Desdín-Micó G, Alonso-Calleja A, Vílchez-Acosta A, et al. Loss of H3K9 trimethylation leads to premature aging. 2024.

36. Oleksiak MF, Rajora OP. Population genomics: Marine organisms: Springer; 2020 2020.

37. Pasyukova EG, Vaiserman AM. HDAC inhibitors: A new promising drug class in anti-aging research. Mechanisms of Ageing and Development. 2017;166:6–15.

38. Patro R, Duggal G, Love MI, Irizarry RA, Kingsford C. Salmon provides fast and bias-aware quantification of transcript expression. Nature Methods. 2017;14(4):417–9.

39. Pawar A, Chowdhury OR, Salvi O. A narrative review of survival analysis in oncology using R. *Cancer Research*, Statistics, and Treatment. 2022;5(3):554–61.

40. Quarni W, Dutta R, Green R, Katiri S, Patel B, Mohapatra SS, et al. Mithramycin A inhibits colorectal cancer growth by targeting cancer stem cells. Scientific Reports. 2019;9(1):15202.

41. Shechter D, Dormann HL, Allis CD, Hake SB. Extraction, purification and analysis of histones. Nature Protocols. 2007;2(6):1445–57.

42. Shimazu T, Hirschey MD, Newman J, He W, Shirakawa K, Le Moan N, et al. Suppression of oxidative stress by β-hydroxybutyrate, an endogenous histone deacetylase inhibitor. Science. 2013;339(6116):211-4.

43. Therneau T, others. A package for survival analysis in S. R package version. 2015;2(7):2014.

44. Vaiserman AM, Kolyada AK, Koshel NM, Simonenko AV, Pasyukova EG. Effect of histone deacetylase inhibitor sodium butyrate on viability and life span in *Drosophila melanogaster*. Advances in Gerontology. 2013;3:30–4.

45. Voisine C, Varma H, Walker N, Bates EA, Stockwell BR, Hart AC. Identification of potential therapeutic drugs for Huntington’s disease using *Caenorhabditis elegans*. PloS one. 2007;2(6):e504.

46. Wang K, Liu H, Hu Q, Wang L, Liu J, Zheng Z, et al. Epigenetic regulation of aging: implications for interventions of aging and diseases. Signal Transduction and Targeted Therapy. 2022;7(1):374.

47. Wang Y, Yuan Q, Xie L. Histone modifications in aging: the underlying mechanisms and implications. Current Stem Cell Research & Therapy. 2018;13(2):125–35.

48. Wickham H, Wickham H. Data analysis. ggplot2: Elegant graphics for data analysis. 2016:189–201.

49. Wickham H, Averick M, Bryan J, Chang W, McGowan LDA, François R, et al. Welcome to the Tidyverse. Journal of Open-Source Software. 2019;4(43):1686.

50. Wickham H, Francois R, Henry L, Müller K. dplyr. A Grammar of Data Manipulation 2020 [Last accessed on 2020 Aug 12] Available from. 2014:Rproject.

51. Wood JG, Hillenmeyer S, Lawrence C, Chang C, Hosier S, Lightfoot W, et al. Chromatin remodeling in the aging genome of *Drosophila*. Aging Cell. 2010;9(6):971–8.

52. Xu D, Fang H, Liu J, Chen Y, Gu Y, Sun G, et al. ChIP-seq assay revealed histone modification H3K9ac involved in heat shock response of the sea cucumber *Apostichopus japonicus*. Science of The Total Environment. 2022;820:153168.

53. Zhao Y, Xuan H, Shen C, Liu P, Han J-DJ, Yu W. Immunosuppression induced by brain-specific HDAC6 knockdown improves aging performance in *Drosophila melanogaster*. Phenomics. 2022;2(3):194–200.

54. Zhao Z, Feng L, Peng X, Ma T, Tong R, Zhong L. Role of histone methyltransferase SETDB1 in regulation of tumourigenesis and immune response. Frontiers in Pharmacology. 2022;13:1073713.

55. Zupkovitz G, Lagger S, Martin D, Steiner M, Hagelkruys A, Seiser C, et al. Histone deacetylase 1 expression is inversely correlated with age in the short-lived fish *Nothobranchius furzeri*. Histochemistry and Cell Biology. 2018;150:255–69.

